# A triad of kicknet sampling, eDNA metabarcoding, and predictive modeling to assess aquatic macroinvertebrate biodiversity

**DOI:** 10.1101/2022.01.03.474789

**Authors:** François Keck, Samuel Hürlemann, Nadine Locher, Christian Stamm, Kristy Deiner, Florian Altermatt

**Affiliations:** Eawag: Swiss Federal Institute of Aquatic Science and Technology, Department of Aquatic Ecology, Überlandstr. 133, CH-8600 Dübendorf, Switzerland; Eawag: Swiss Federal Institute of Aquatic Science and Technology, Department of Environmental Chemistry, Überlandstr. 133, CH-8600 Dübendorf, Switzerland; ETH, Department of Environmental Systems Science, Universitätstr. 16, CH-8092 Zürich, Switzerland; Department of Evolutionary Biology and Environmental Studies, University of Zurich, Winterthurerstr. 190, CH-8057 Zürich, Switzerland

**Keywords:** Metabarcoding, water DNA, Ephemeroptera, Plecoptera, Trichoptera

## Abstract

Monitoring freshwater biodiversity is essential to understand the impacts of human activities and for effective management of ecosystems. Thereby, biodiversity can be assessed through direct collection of targeted organisms, through indirect evidence of their presence (e.g. signs, environmental DNA, camera trap, etc.), or through extrapolations from species distribution models (SDM). Differences in approaches used in biodiversity assessment, however, may come with individual challenges and hinder cross-study comparability. In the context of rapidly developing techniques, we compared a triad of approaches in order to understand assessment of aquatic macroinvertebrate biodiversity. Specifically, we compared the community composition and species richness of three orders of aquatic macroinvertebrates (mayflies, stoneflies, and caddisflies, hereafter EPT) obtained via eDNA metabarcoding and via traditional *in situ* kicknet sampling to catchment-level based predictions of a species distribution model. We used kicknet data from 24 sites in Switzerland and compared taxonomic lists to those obtained using eDNA amplified with two different primer sets. Richness detected by these methods was compared to the independent predictions made by a statistical species distribution model using landscape-level features to estimate EPT diversity. Despite the ability of eDNA to consistently detect some EPT species found by traditional sampling, we found important discrepancies in community composition between the two approaches, particularly at local scale. Overall, the more specific set of primers, namely fwhF2/EPTDr2n, was most efficient for the detection of target species and for characterizing the diversity of EPT. Moreover, we found that the species richness measured by eDNA was poorly correlated to the richness measured by kicknet sampling and that the richness estimated by eDNA and kicknet were poorly correlated with the prediction of the statistical model. Overall, however, neither eDNA nor the traditional approach had strong links to the predictive models, indicating inherent limitations in upscaling species richness estimates. Future challenges include improving the accuracy and sensitivity of each approach individually yet also acknowledge their respective limitations, in order to best meet stakeholder demands addressing the biodiversity crisis we are facing.

## Introduction

The role of biodiversity in maintaining ecosystem functions and services is widely recognized (Chapin 2000, Cardinale 2012). Consequently, deleterious effects of human activities on biodiversity are a source of growing concern and are mobilising both scientists and stakeholders around the world (Pereira & Cooper 2006, Diaz et al. 2020). In a context where the loss of biodiversity is established and threatens many of the benefits that ecosystems provide to humanity, monitoring the diversity and composition of biological communities is a priority, both to prevent future adverse consequences and to establish possible restoration measures (Lindenmayer & Likens 2010). However, measuring state and change of biodiversity remains a challenge both due to questions related to its scientific definition (such as which levels of biological organisation to study and at what spatial scales) and to the limitation of the methods and technologies available to monitor life in the environment.

For a long time, freshwater biodiversity monitoring has solely relied on the capture of individuals or their direct observation. These approaches, although improved over time, remain limited by sampling biases, identification errors, associated costs, and sometimes coarse taxonomic resolution. Furthermore, they do not allow upscaling and predicting to larger spatial or temporal scales. Thus, additional approaches are needed to complement classic biodiversity data, especially with respect to a better scaling and resolving the state and change of biodiversity. Approaches can be based on novel technological advances, such as in molecular sciences, or in a more detailed use of predictive or other statistical models (Guisan and Zimmermann 2000; Taberlet et al. 2012; Petchey et al. 2015; Altermatt et al. 2020). The implementation of these approaches, however, needs to be complemented with a thorough analysis of strengths and weaknesses, including directly comparing performance of the approaches as well as identifying what can (or cannot) be gained by either approach. Within the last decade, environmental DNA (eDNA) has been – especially in aquatic ecosystems – presented as a game-changer to traditional approaches, with the promise of being able to monitor biodiversity at unprecedented spatial and temporal scales (Hering et al., 2018; Leese et al., 2016, Deiner et al. 2017). In streams and rivers, it has also already been extensively used and compared to classic kicknet-based approaches, and complementarity and respective advantages and disadvantages have been put forward (e.g. Mächler et al., 2019, Hänfling et al., 2016, Pont et al. 2018). Several recent meta-analyses (Keck et al. 2021; McElroy et al. 2020) showed that, in aquatic environments, eDNA metabarcoding and traditional methods can provide similar estimates of taxonomic richness, but large inconsistencies remain in the taxonomic composition found by the two approaches, especially in macroinvertebrate and microbial communities.

A pairwise comparison of methods, however, may be hard to resolve, as either method could be a better approximation of reality. Thus, including a third approach, using a triad of comparisons (Figure 1), offers the possibility to resolve such discussions, yet hinges on models that rely on independent and exogenous variables (e.g. environmental variables) to predict diversity (see e.g. Moraes et al. 2014; Lobo et al. 2004; Lehmann et al. 2002). This latter approach does not estimate diversity from direct observation but from mathematical functions or statistical relationships previously established (Ferrier and Guisan 2006). Since direct observations (traditional or DNA-based) are still very sparse and limited, this third approach is the only one that currently allows us to estimate biodiversity on a large scale and in a continuous manner. However, there has been little – if any – work on linking the estimates obtained by such models (usually trained with traditional observational data) with those obtained from eDNA.

**Figure 1:**
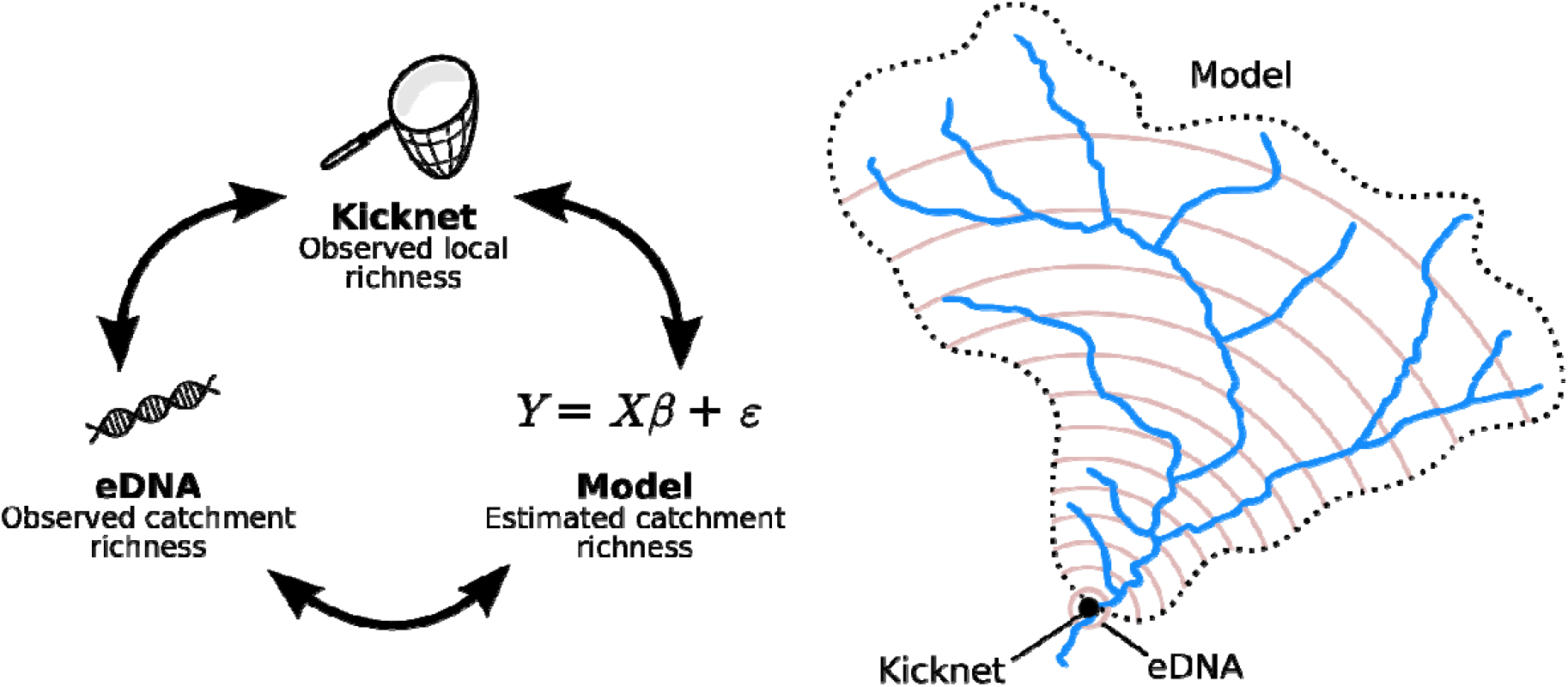
A triad of methods (kicknet sampling, eDNA sampling, and statistical modelling) available to estimate macroinvertebrate diversity in river ecosystems. Each has its own specificities, particularly in terms of integrated spatial scale. Note that models always rely on underlying data used to train them, in this study those are independent kick-net samples.

In this study, we used a dataset of 24 streams located in Switzerland, for which macroinvertebrate communities have been sampled at one location, both by kicknet and eDNA, and for which independent predictions on species richness have been modelled. We specifically focus on the diversity of three orders of macroinvertebrates: mayflies (Ephemeroptera, E), stoneflies (Plecoptera, P), and caddisflies (Trichoptera, T). EPT taxa are commonly found in streams and rivers, and have proven to be useful and powerful indicators of water quality (Wallace et al. 1996). We amplified eDNA with two distinct pairs of primers, a more generic one (mlCOIintF/HCO2198, Leray et al. 2013, Folmer et al. 1994) and one more specific toward benthic invertebrate taxa (fwhF2/EPTDr2n, Vamos et al. 2017, Leese et al. 2021), in order to test their respective capacity to unveil EPT diversity. We compared the diversity estimates and the species composition detected by the eDNA and kicknet approaches, both at regional (gamma diversity) and local (alpha diversity) scale. We then related these results to the diversity estimated by a predictive statistical model for EPT richness (Kaelin and Altermatt 2016). Our goal was to evaluate the ability of this triad of methods to estimate and characterize the biodiversity in streams, and to investigate their differences.

## Material and Methods

### Sampling

Water samples were collected from 24 streams in Switzerland in 2013–2014 (Figure 2). All streams were small to medium sized streams (range of catchment area 7 to 66 km^2^) in the Plateau and Jura part of Switzerland, covering an elevational range from 370 to 912 m a.s.l. All were headwater streams with no waste water treatment plants upstreams, and land-use types in the upstream catchment consisted mostly of forest and agriculture (dairy farming and cropping). Settlements covered between 5 and 21% of the catchment areas. At each location, we sampled two sites in the stream located a few hundreds meters apart, yet within the same habitat type and environmental conditions. Macroinvertebrate communities were sampled using kicknets and water samples were collected for eDNA analyses. Water samples were transported in a cooler on ice (maximum transport time of six hours) and were stored at –20 °C until processed further. All samples were taken within a larger research program (for details of the project and sampling procedure, see also Burdon et al. 2019, Stamm et al. 2016, 2017). Here we focus on the subset of samples taken upstream of waste water treatment plant inflows only.

**Figure 2:**
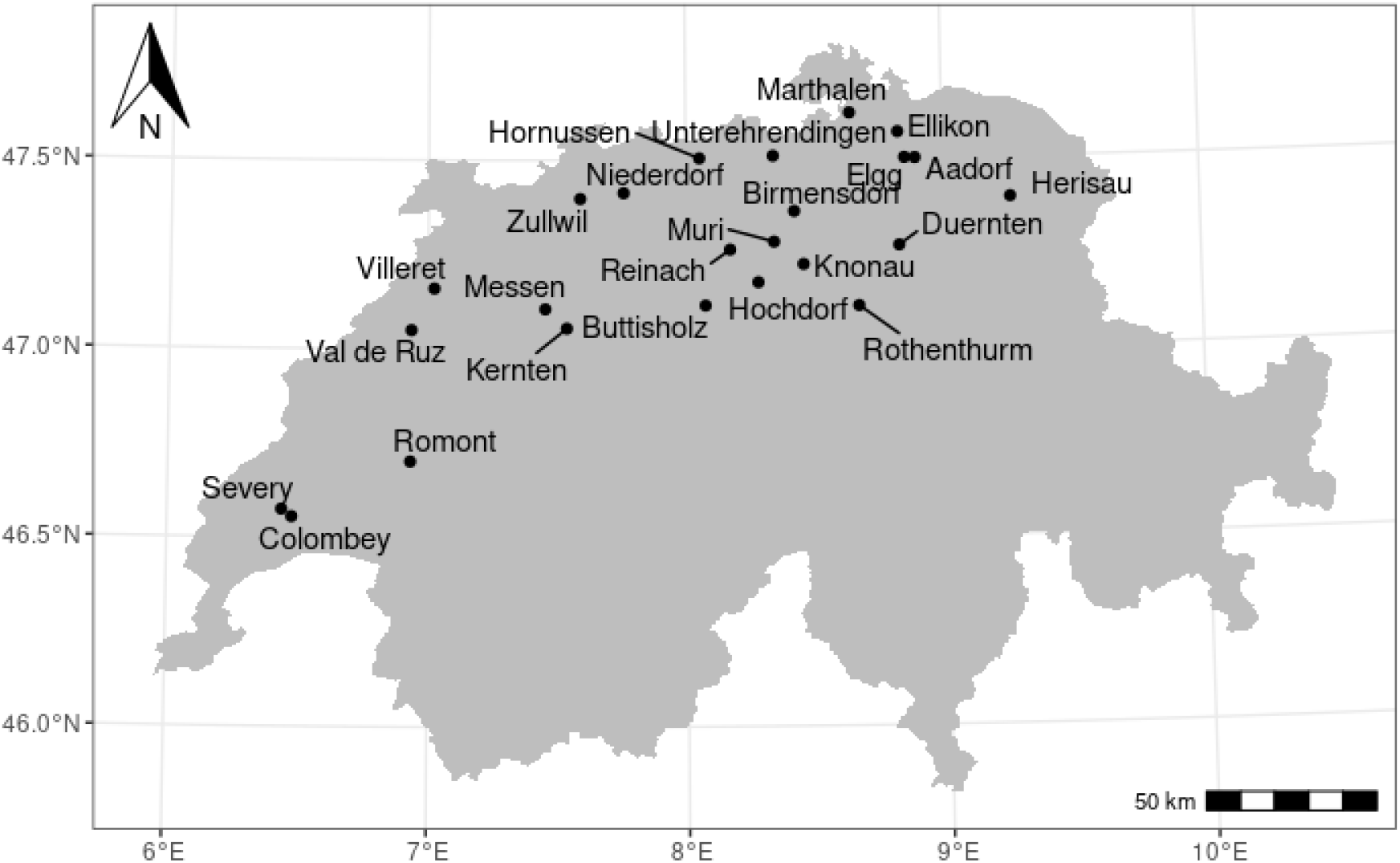
Map of Switzerland showing the 24 sampling locations. Locations are named after local municipalities.

### EPT identification

At each location, all individuals of may-, stone-, and caddisflies (EPT) were identified to the species level (in few cases to pre-defined species complexes, subsequently treated as species) using expert taxonomists. Identification of all taxa followed pre-defined taxonomic lists, and all data from the two sites per location were pooled. For details see Burdon et al. (2019) and Stucki (2010). For subsequent analyses, we only used presence/absence data, and calculated species richness values per location.

### Water filtration and DNA extraction

Methods for filtration and extraction of DNA from water samples were previously published in Mansfeldt et al. (2020). Briefly, water was filtered through a glass fiber filter (GF/F, nominal pore size of 0.7 µm, 25 mm, Whatman International Ltd., England) and was extracted with a Phenol-Chloroform Isoamyl followed by an ethanol precipitation (Mansfeldt et al. 2020). Strict adherence to contamination control was followed using a controlled lab where only eDNA isolation and pre-PCR preparations are performed (Deiner et al. 2015). Between two and eight independent extractions from filters were carried out for each sample location. Total volume of water filtered for each extraction depended on the suspended solids in the sample, which clogged the filter, and ranged from 65 to 350 mL. A total of 500 to 700 mL of filtered water was used per sample for DNA extraction (see Mansfeldt et al. 2020). A 50 µL pool was created by adding equal volumes from each independent extraction and quantified using the Qubit (1.0) fluorometer following recommended protocols for the dsDNA HS Assay, which has a high accuracy for double stranded DNA between 1 ng/mL to 500 ng/mL (Life Technologies, Carlsbad, CA, USA). Filter negative controls were created for each day that filtration took place. A filter negative control consisted of filtering 250 mL of Milli-Q® water that was secondarily decontaminated with UVC light. DNA extraction controls were used to monitor contamination and processed with each batch of extractions of which consisted of between 18 and 22 filters per batch (Table SXX: Controls tab). All pooled DNA extractions were cleaned with the OneStepTM PCR Inhibitor Removal Kit (Zymo Research, Irvine, California, USA) according to the manufacturer’s protocol as this has been shown to be effective for removal of PCR inhibition of riverine samples of environmental DNA (McKee et al. 2015).

### Library construction and sequencing

Library construction for each sample location followed a three step PCR process. The first PCR consisted of amplification of a 312 bp fragment of the 5’ end of the Cytochrom Oxidase I mitochondrial gene (COI) using the forward primer (mlCOIintF) from Leray et al. (2013) and the reverse primer (HCO2198) from Folmer et al. (1994). Four independent PCRs on eDNA were carried out in 15 µL volumes with final concentrations of 1x supplied buffer (Faststart TAQ, Roche, Inc., Basel, Switzerland), 1000 ng/µL BSA (New England Biolabs, Ipswich, MA, USA), 0.2 mMol dNTPs, 2.0 mMol MgCl2, 0.05 units per µL Taq DNA polymerase (Faststart TAQ, Roche, Inc., Basel, Switzerland), and 0.5 µMol of each forward and reverse primer. 2 µL of extracted eDNA was added that ranged in concentration from 0.03 to 54.0 ng/µL. This range was the outcome of DNA concentrations that were extracted. The thermal-cycling regime was 95 °C for 4 minutes, followed by 35 cycles of 95 °C for 30 seconds, 48 °C for 30 seconds and 72 °C for 1 minute. A final extension of 72 °C for 5 minutes was carried out and the PCR was cooled to 4 °C until removed and stored at –20 °C until products were cleaned. PCR products were visualized on a 1.5% agarose gel to confirm amplification. We cleaned each PCR replicate with Exo I Nuclease (EXO I) and Shrimp Alkaline Phosphatase (SAP) (Thermo Fisher Scientific Inc., Waltham, Maryland USA). The master mix consisted of 1.6 U/µL Exo I and 0.15 U/µL SAP in a total volume of 1.1 µL which was then added to 7.5 µL of the PCR product. Products were heated to 37 °C for 15 minutes and followed by 15 minutes at 80 °C for deactivation of EXO and SAP.

The second PCR was conducted with the same PCR conditions above except the forward and reverse primers were modified to include the Nextera® transposase adaptors and only 1 µL of cleaned PCR product was used in the reaction. Between the forward and reverse primer sequence and the transposase adaptor a different number of random bases were inserted to create products of varying length to allow more heterogeneity on the flow cell. The thermal-cycling regime was the same except that five cycles were used. PCR products from the four independent reactions for each sample were then pooled together and cleaned using a two-step method. First, we cleaned each pooled reaction with EXO I and SAP as described above except we adjusted proportionally the volumes of EXO I and SAP for a total cleaned volume of 30 µL rather than 7.5 µL. Second, we desalted, removed buffer components with the Illustra MicroSpin S-300 HR Columns (GE Healthcare Life Sciences, Little Chalfont, United Kingdom) following the manufacturer’s recommended protocol.

The third PCR was to index each pooled PCR by before pooling all PCR from each site for sequencing. We duel-indexed samples using the Nextera® index kits A and D. PCR was carried out in 50 µL were samples were added at either 5 or 10 µL, where amplicons that showed a DNA concentration less than 0.1 ng/µL were added at 10 µL and all other greater than this were added at 5 µL. We used the KAPA Library Amplification Kit following the manufacturer’s recommended protocol (KAPA Biosystems, Wilmington, MA). Each of the pooled reactions were then cleaned using Agencourt AMPure XP beads following the recommended manufacturer’s protocol (Beckman Coulter, Brea, CA, USA).

Cleaned and indexed libraries were then assayed for DNA concentration using the Qubit (1.0) fluorometer following recommended protocols for the dsDNA HS Assay, normalized then pooled at a 2 nM concentration. PHiX control was added at 1%. Paired-end sequencing was performed on an Illumina MiSeq (MiSeq Reagent kit v2, 250 cycles) at the Genomic Diversity Center at the ETH, Zurich, Switzerland following manufacturer’s run protocols (Illumina, Inc., San Diego, CA, USA). The MiSeq Control Software Version 2.2 including MiSeq Reporter 2.2 was used for the primary analysis and the demultiplexing of the raw reads.

In order to amplify the 142 bp long fragment of the COI locus using fwhF2 forward primer (Vamos et al. 2017) and EPTDr2n reverse primer (Leese et al. 2021) a similar three-step PCR as described above, was conducted. First PCR was carried out in three independent PCR reactions with a total volume of 25 µL containing final concentrations of 1x supplied buffer (Faststart TAQ, Roche, Inc., Basel, Switzerland), 1500 ng/µL BSA (Molecular biology grade, New England Biolabs), 0.2 mMol dNTPs, 3.0 mMol MgCl2, 0.05 units per µL Taq DNA polymerase (Faststart TAQ, Roche, Inc., Basel, Switzerland), and 0.5 µMol of each forward and reverse primer. 2 µL of extracted eDNA or PCR grade water as negative control was added to each reaction. PCR Reactions were performed with the following cycle settings on a (Biometra T1Thermocycler, Analytik Jena GMBH, Ge): denaturation was at 95°C for 8 minutes, followed by 30 cycles of 95 °C for 30 seconds, 50 °C for 1 minute and 72°C for 1 minute. A final extension of 72 °C for 7 minutes was performed, followed by lowering the temperature to 4°C to avoid DNA degrading.

From the first PCR product, 10 µL was enzymatically cleaned by adding 0.11 U/µL Exonuclease I (E. coli), 0.2 U/µL Shrimp Alkaline Phosphatase (rSAP) (New England Biolabs) and 1.11 µL PCR grade water to each sample. The temperature cycling was carried out, as recommended by the manufacturer.

In order to add the Nextera transposase sequences adaptors to the first PCR fragment, 4 µL cleaned PCR product was used in similar PCR condition as in the first PCR reaction. Thermal cycling regime was identical, except that the number of cycles were reduced. Amplification success was checked with the AM320 method on the QiAxcel Screening Cartridge (Qiagen, Germany). Most of the samples worked after 10 PCR cycles. However, the cycling number for 28 samples was adjusted up to 18 cycles, in order to see amplification success.

Before we attached the index adapters with the third PCR, additional cleaning steps were performed. This consisted of first pooling the replicates of the second PCR product and then running it on a 0.8% low melting point Agarose (Analytical grade, Promega) together with 100-bp ladders (Promega, Madison, WI, USA). Fragments with the correct size of 268 bp were cutted out from gel, by using a fresh scalpel. Thereafter DNA was purified, using the Wizard SV Gel and PCR Clean-Up System (Promega, Madison, WI, USA). Exciseds DNA bands were dissolved in 250 µL Membrane Binding Solution at 65 °C shaken at 850 rpm for 2 minutes. After the column bind and washing steps, DNA was eluted in 20 µL PCR grade water.

Illumina Nextera XT Index set D (Illumina, Inc., San Diego, CA, USA) were attached to the purified amplicon by following the recommended protocol from the Illumina library preparation guide, except increasing cycle number from 8 to 10 cycles. After the Nextera® index adapters successfully bound to the fragment, the individual samples were cleaned up with a MagJET NGS Cleanup and Size Selection Kit running on a KingFisher Flex Purification System (Thermo Fisher Scientific Inc., MA, USA).

Quantification of PCR products was conducted with a target selective fluorescence dye Qubit BR DNA Assay Kit (Life Technologies, Carlsbad, CA, USA). Fluorescence dye emission of the standard dilution series and samples were measured in replicates with a Spark Multimode Microplate Reader (Tecan, US Inc., USA). Samples, including filter, extraction and PCR controls were then merged in four equimolar pools (3nM), in relation to their concentration, with an automated liquid handling station (BRAND GMBH + CO KG, Wertheim, GE). Final pool was then three times manually purified, by using a 0.8x ratio of Agencourt AMPure XP (Beckman Coulter, Brea, CA, USA) beads, again following the recommended manufacturer’s protocol. Amplicon size was verified by an Agilent 4200 TapeStation (AgilentTechnologies, Inc., USA) run. Library was sequenced with a concentration of 10 pM in the flowcell on an Illumina MiSeq (Illumina, Inc. San Diego, CA, USA) at the Genetic Diversity Center (ETH, Zurich). The Sequencing run (MiSeq Reagent kit v2, 300 cycles, paired-ended) was spiked with 10% PHiX control.

### Bioinformatics

The software package DADA2 v.1.16.0 was used to infer amplicon sequence variants (ASVs) from the demultiplexed MiSeq (forward and reverse) reads following the methods described by Callahan et al. (2016). Primer sequences (mlCOIintF/HCO2198 and fwhF2/EPTDr2n) were removed from the reads using cutadapt v.2.10 (Martin, 2011). After primer removal, the forward and reverse reads were truncated to 200 and 170 nucleotides respectively for the mlCOIintF/HCO2198 run, in order to remove poor quality nucleotides at their extremities. Both the forward and reverse reads were truncated to 120 nucleotides for the fwhF2/EPTDr2n run. Reads were quality-filtered by removing any read with one or more ambiguities (“N”) and any read with a maximum expected error (maxEE) larger than 2. After dereplication, ASVs were finally selected based on the error rates model determined by the DADA2 denoising algorithm and paired reads merged into one sequence using a minimum overlap of 12 bases. Potential chimeric sequences were removed using the de novo bimera detection algorithm implemented in DADA2.

We translated the ASV sequences into amino acids starting from the 2nd nucleotide and using the invertebrate mitochondrial code. Since COI is a coding sequence, it is not expected to find stop codons in the barcode region. Therefore, all the ASV sequences (2642 for the mlCOIintF/HCO2198 primers, 2251 for the fwhF2/EPTDr2n primers) in which a stop codon was found were discarded. For the mlCOIintF/HCO2198 run, a total of 140 additional ASVs which were found in relative proportion > 0.1% in one of the six negative controls were also discarded from all the samples. For the fwhF2/EPTDr2n run, only 2 ASV sequences were removed at this step (2 negative controls were used).

Taxonomic assignment of ASV sequences was achieved using the RDP algorithm (Wang et al. 2007) with a bootstrap threshold of 75%. The reference database used for taxonomic assignment was assembled from several sources: NCBI, Bold, MIDORI and the EPT sequences collected within the SwissBOL project. After quality filtering (removing incorrect sequences and mislabeled taxa) the reference database included 654,132 labeled COI sequences divided in 88 classes, 493 orders, 4,107 families, 33,337 genera and 120,374 species. Replicates (sites) were merged by locations. For five locations (Buttisholz, Hochdorf, Hornussen, Messen, and Niederdorf, see Figure 2), only one replicate was available for mlCOIintF/HCO2198. Therefore we excluded the corresponding replicates from the analysis of fwhF2/EPTDr2n.

### Predictive model for EPT richness

For each sampling location, we predicted the EPT species richness using a statistical species distribution (species richness) model developed by Kaelin and Altermatt (2016), and model predictions were directly taken from that publication for the respective 24 study catchments used here. Briefly, this model is a generalized linear model using a Poisson error distribution. The model was trained to predict EPT species richness from a set of 11 environmental variables using lasso regularization. The model had been trained with a dataset of 410 independent locations where EPT species richness was assessed by kicknet sampling. These 410 locations did not overlap with any of the 24 study locations/catchments herein used, and had been monitored by kicknet in a systematic manner between 2009– 2013, ensuring random spatial and temporal coverage (for details, see Altermatt et al. 2013, Ryo et al. 2018). These sites cover a much wider environmental, geographic and temporal scale than the 24 study catchments compared to, thus should encapsulate all variation in species richness expected in the latter. Then, using generalized linear models incorporating all main land-use variables identified as relevant by Kaelin & Altermatt (2016), the model was used to predict species richness in 22,169 ∼2 km^2^ large sub-catchments, covering the entire territory of Switzerland. Predictions on alpha diversity (richness) of EPT were retrieved for the sub-catchments corresponding to the 24 locations studied here. Thus, the predictive species distribution model made predictions on the expected richness in the 24 study catchments further analysed here are based on a model parametrized across all of Switzerland. We note that the data used to train the predictive model are also based on kicknet samples. That is, there may be an inherent part of diversity only detectable by eDNA that cannot be assessed by the kicknet method, which would thus also not be covered by the model. Importantly, however, the model makes only predictions at the level of total richness, and not at the level of individual species’ identity. Thus, predictions are at a coarser level, such that this effect is not expected to play a major role, or maximally result in a shift in the intercept of richness predictions.

### Analyses

We used presence-absence data and species richness (i.e. the number of species) to characterize the diversity of EPT, both from the eDNA as well as the kicknet data. Diversity was studied both at local scale (i.e. locations after merging site replicates, alpha diversity), and at regional scale (i.e. all locations merged, gamma diversity). For both alpha and gamma diversity, we compared the number of species detected by kicknet only, by eDNA only, and by the two approaches simultaneously. For each location, the sampling effort (number of identified individuals and sequencing depth) was assessed with species accumulation curves. Finally, we computed and tested Pearson correlations between the richness found by eDNA (fwhF2/EPTDr2n and mlCOIintF/HCO2198 primers separately), found by kicknet and estimated by the predictive model. Analyses were conducted using R 4.0.3 (R Core Team, 2020).

### Data and code

All raw sequencing data are available at the European Nucleotide Archive (ENA) under the accession number PRJEB26649. The processed data and R scripts to reproduce the analyses and results are available at : https://github.com/fkeck/ecoimpact.

## Results

Library sequencing generated 4,638,809 sequences (mlCOIintF/HCO2198 primers) and 8,008,677 sequences (fwhF2/EPTDr2n primers). For sequences amplified using the mlCOIintF/HCO2198 primers, the pre-processed and quality-filtered data consists of 3,110,057 reads divided in 13,797 ASVs. For sequences amplified using the fwhF2/EPTDr2n primers, the pre-processed and quality-filtered data consists of 4,779,863 reads divided in 2,665 ASVs.

For the mlCOIintF/HCO2198 primers, taxonomic assignment failed for a significant number of ASVs for which identification was not possible, even at the highest taxonomic ranks (87% of unclassified Eukaryota). Assigned reads are dominated by insects (Diptera, Coleoptera and unclassified Insecta), Clitellata, Chromadorea and unclassified arthropods. The orders of interest (EPT) only represent a small proportion of assigned ASVs (7%), with 32 Ephemeroptera, 17 Plecoptera and 34 Trichoptera taxa detected. The relative proportion of EPT is even less important when accounting for the number of reads. In total the EPT groups represent 3.1% of the assigned reads. In contrast, the fwhF2/EPTDr2n primers performed better with a lower proportion of unidentified Eukaryota (47.9%). Targeted orders were also more represented with 63 ASVs identified as Ephemeroptera, 37 as Plecoptera, and 42 as Trichoptera taxa, representing 10% of the assigned ASVs (8.6% of the assigned reads). The sampling depth (number of reads identified as EPT) was highly variable among locations (ranging from 7 at Aadorf with mlCOIintF/HCO2198 to 109,956 at Zullwil with fwhF2/EPTDr2n). The absolute number of reads identified as EPT was 10 to 100 times higher with the fwhF2/EPTDr2n primers than with the mlCOIintF/HCO2198 primers (Supplementary Information Figure 1 and 2). In one location (Hornussen) none of the tested primers could detect EPT taxa. However, all the species accumulation curves seem to reach a plateau in the other locations (Supplementary Information Figure 1 and 2). This was not the case with the kicknet data (Supplementary Information Figure 3).

**Figure 3:**
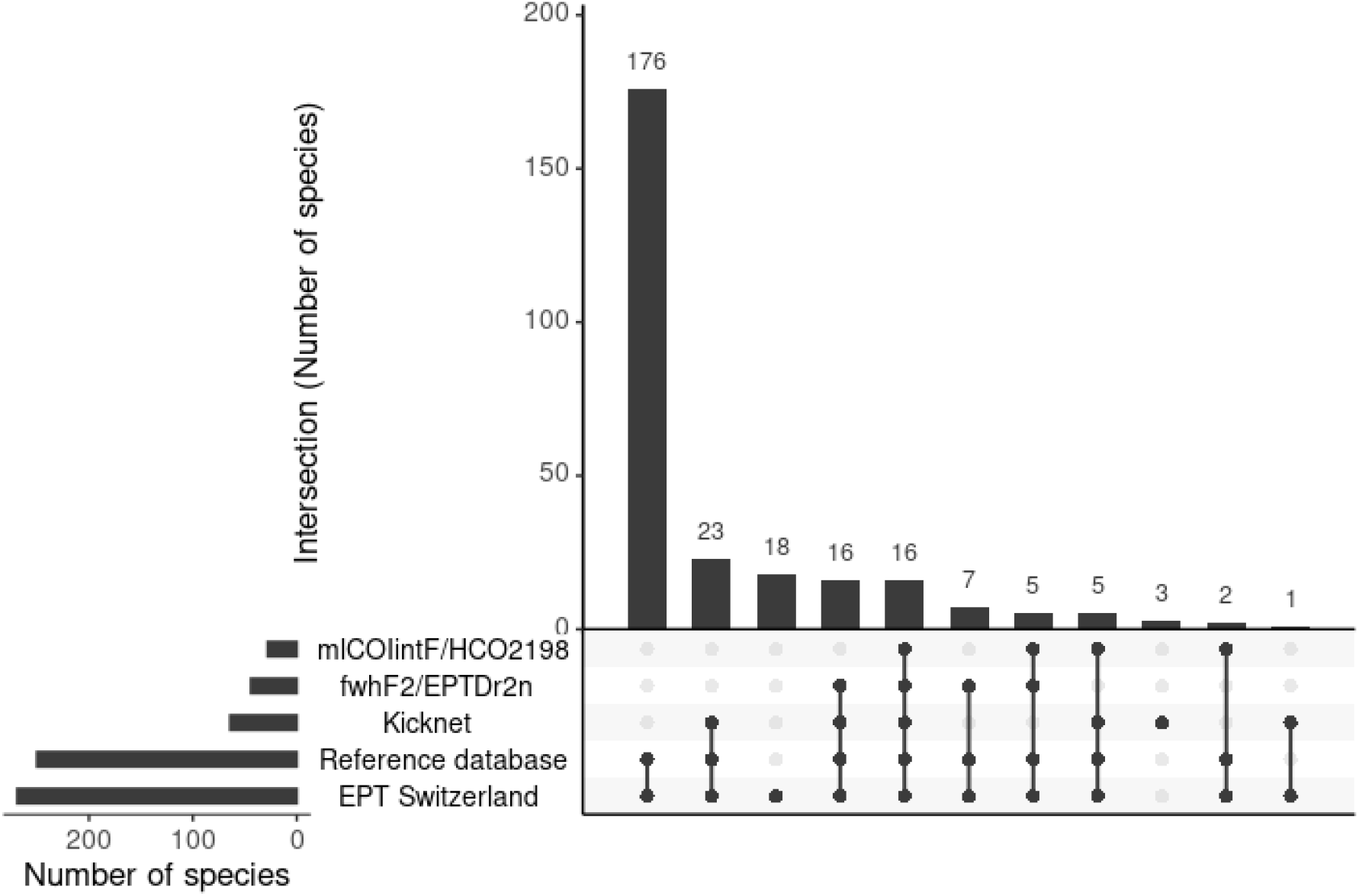
Regional EPT species richness (diversity across all sampling locations) detected by eDNA (mlCOIintF/HCO2198 and fwhF2/EPTDr2n primers) and kicknet method in comparison to total EPT richness known from Switzerland and the subset of species included in the molecular reference database. Horizontal bars show the total number of species in each set. The vertical bars show the number of species in each intersection between sets.

Across all sites (i.e., gamma diversity), kicknet was the method that detected the highest number of different EPT taxa (64), followed by eDNA amplified with the fwhF2/EPTDr2n primers (44 taxa). Results of the regional EPT species richness (across all locations) are shown on Figure 3. Environmental DNA amplified by the mlCOIintF/HCO2198 primers detected only 28 taxa across all sites. In total, 16 taxa were detected by the three methods. We found a better congruence between the fwhF2/EPTDr2n primers and the kicknet (32 common taxa) than between the mlCOIintF/HCO2198 primers and the kicknet (21 common taxa), or between the two primers (21 common taxa).

The number of EPT taxa detected varied both across locations and methods (Figure 4). Additionally, the mlCOIintF/HCO2198 primers did not detect any EPT taxa in three other locations (Buttisholz, Knonau and Rothenthurm). Some locations showed particularly poor diversity (e.g. Colombey, Val de Ruz), while others exhibited a high EPT richness (e.g. Rothenthurm when assessed with the fwhF2/EPTDr2n primers). Overall, alpha diversity (local species richness) was higher with kicknet (mean = 19.6, sd = 6.5) than with eDNA amplified with mlCOIintF/HCO2198 primers (mean = 4.37, sd = 3.85) or fwhF2/EPTDr2n primers (mean = 7, sd = 7.88). The mean richness detected by the fwhF2/EPTDr2n primers was not significantly higher than the mean richness detected by the mlCOIintF/HCO2198 primers (paired t-test, t = -1.48, p-value = 0.15).

**Figure 4:**
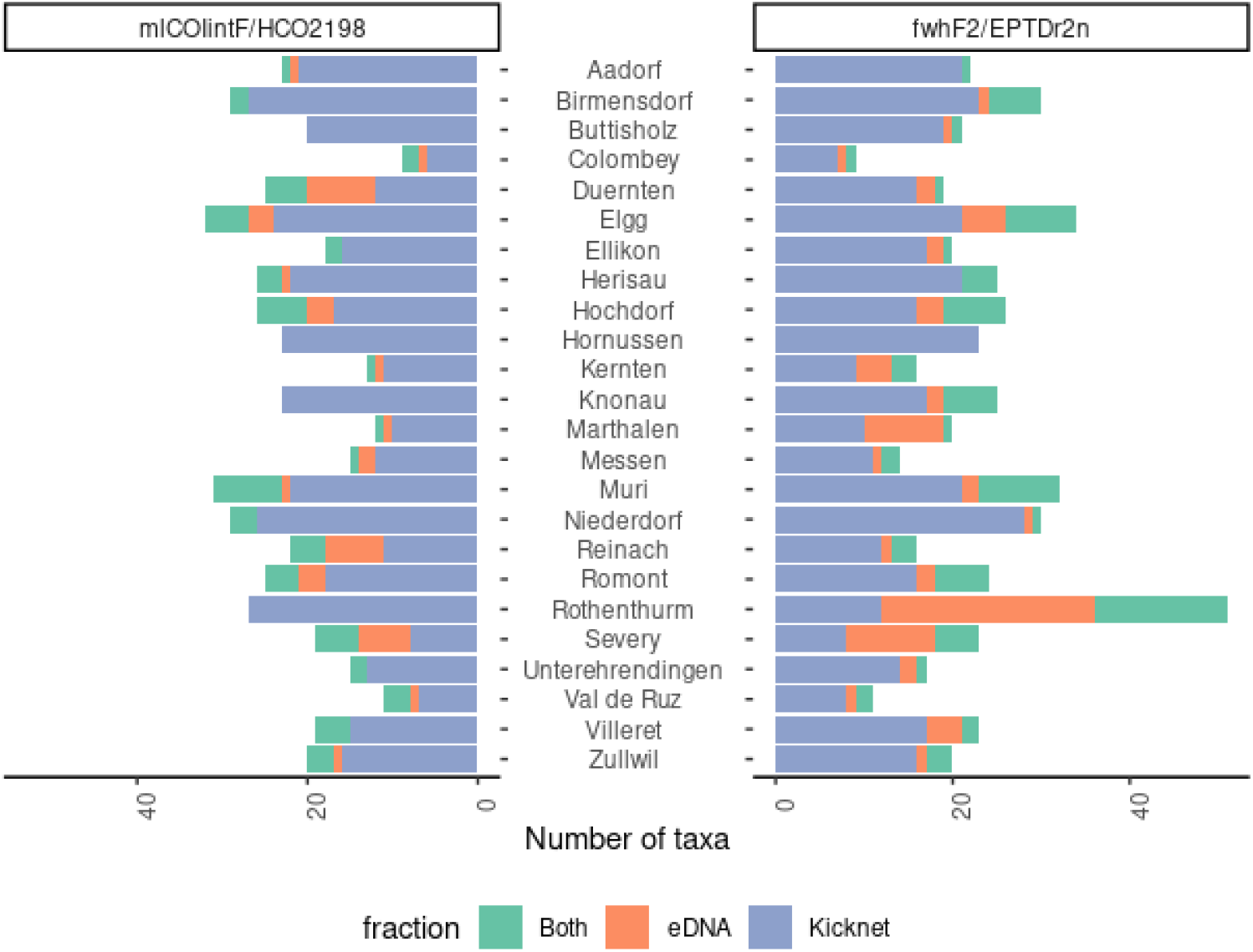
Number of EPT taxa detected in each location by eDNA (mlCOIintF/HCO2198 and fwhF2/EPTDr2n primers) and kicknet methods. The total number of taxa detected is divided in three fractions (in green the taxa detected by the two methods, in orange the taxa detected by eDNA only, and in blue the taxa detected by kicknet only, respectively).

Some taxa commonly detected by kicknet sampling were never or rarely detected by eDNA (Figure 5). For example, this is the case for *Alainites muticus, Centroptilum luteolum, Habrophlebia lauta* or the genus *Hydropsyche*. Contrastingly, the very common species *Baetis rhodani* was well detected by both approaches. There is no common species detected systematically by eDNA that is not detected by the traditional sampling. However, a few species were detected only by eDNA in a few streams (e.g. *Glyphotaelius pellucidus, Nemurella pictetii*, and the *Hydroptila*-complex).

**Figure 5:**
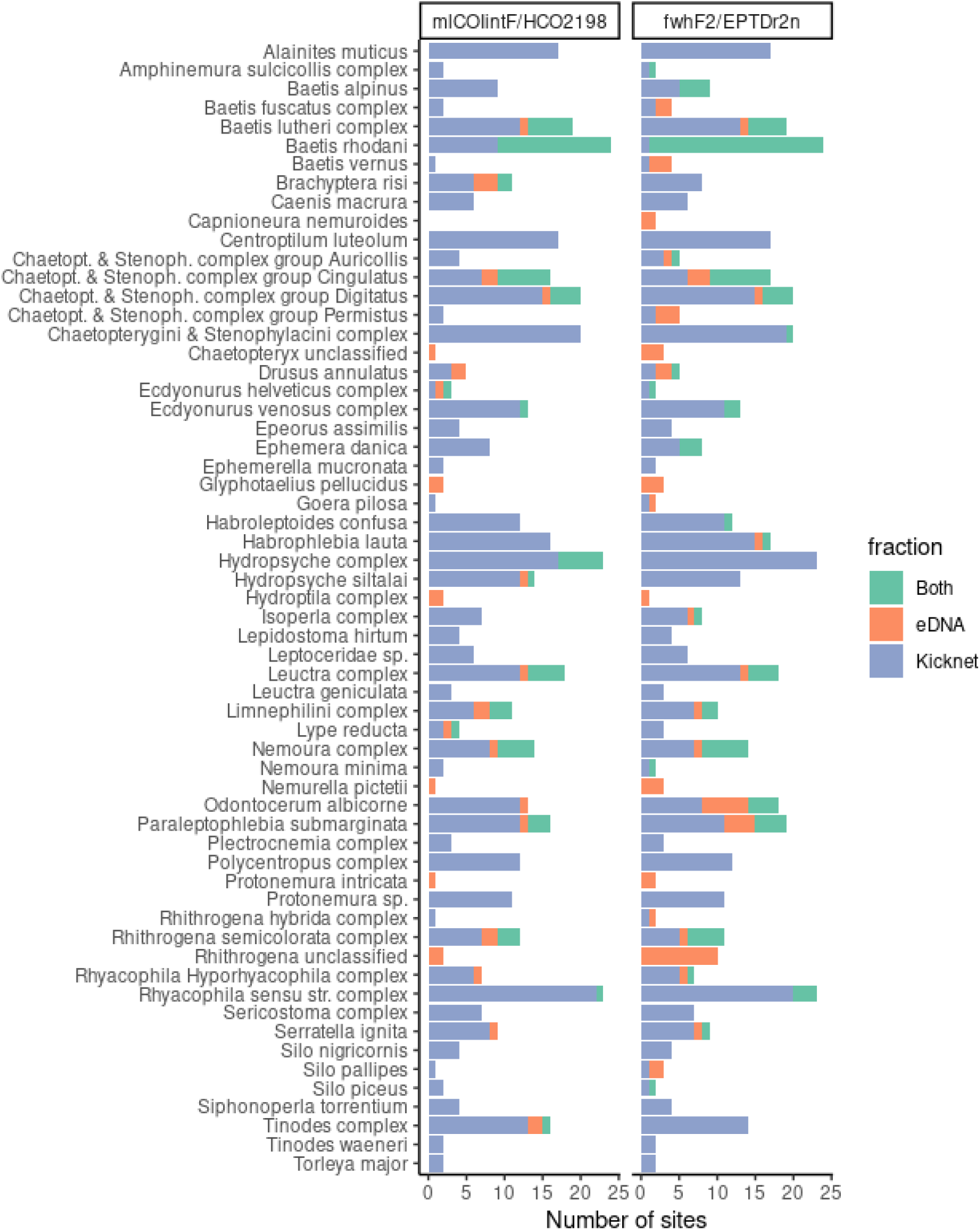
Number of streams where each EPT taxon was detected by eDNA (mlCOIintF/HCO2198 and fwhF2/EPTDr2n primers) and kicknet methods. The total number of locations is divided in three fractions (in green the locations where the taxon was detected by the two methods, in orange the locations where the taxon was detected by eDNA only, and in blue by kicknet only, respectively). For clarity, only the taxa detected more than once (all streams and methods combined) are shown.

We found the correlation between the richness estimates provided by the different methods to be remarkably low (Figure 6). The highest correlation (rho = 0.44, p-value = 0.03) was found between the predictive model and eDNA amplified with the fwhF2/EPTDr2n primers. Correlations between the kicknet method and the predictive model (rho = 0.3, p-value = 0.16) and between the kicknet method and the fwhF2/EPTDr2n primers (rho = 0.27, p-value = 0.2) were not significant. The correlations between the mlCOIintF/HCO2198 primers and the other approaches were close to zero and non-significant (Figure 6). Merging the primers did not improve the correlations between the richness found by eDNA and the other methods (Supplementary Information Figure 4).

**Figure 6:**
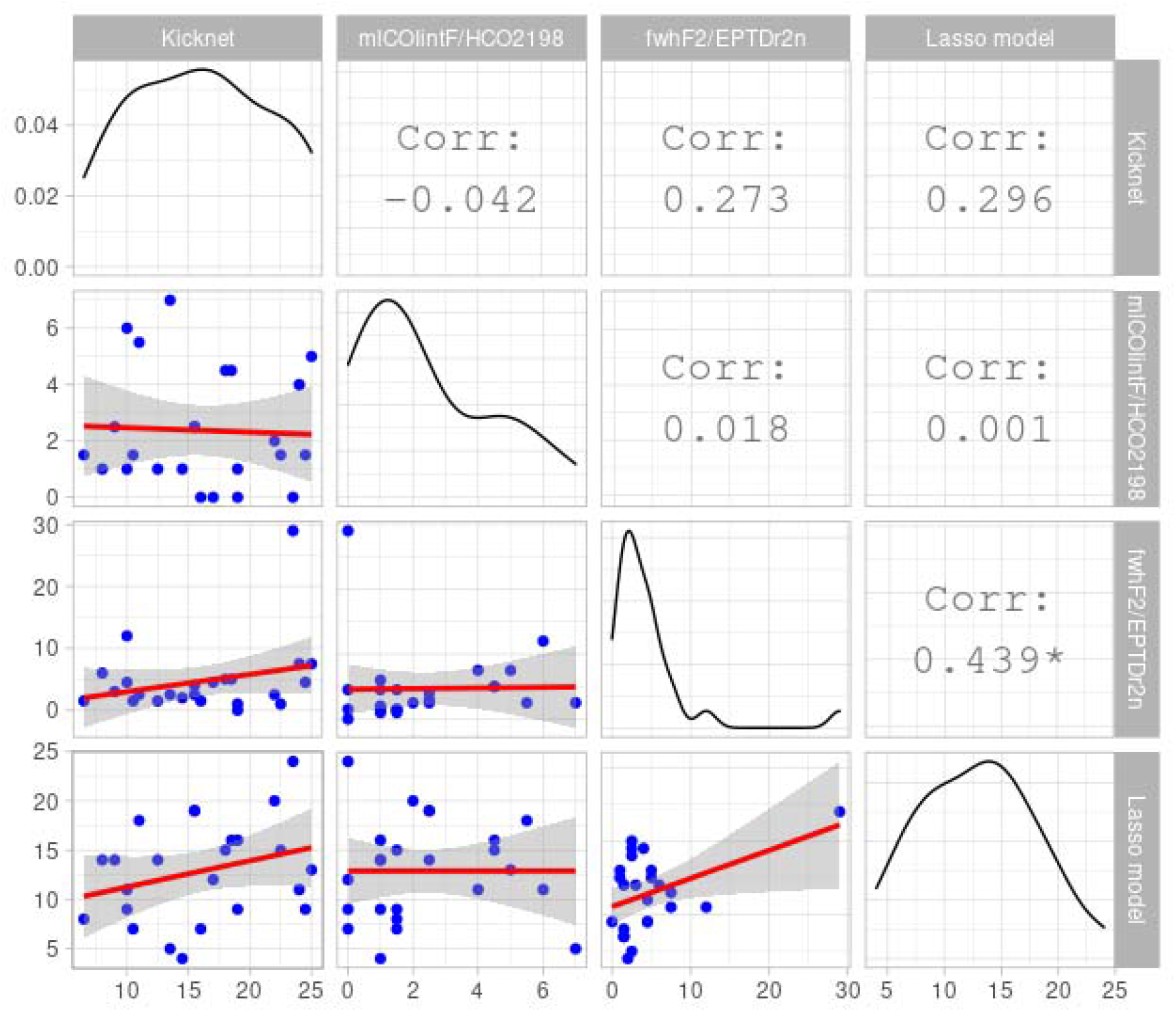
Relationships between the EPT richness estimates provided by the four investigated methods. The upper triangle provides the correlation values between each method (star indicates p-value < 0.05). Lower triangle shows the scatterplots with linear regressions (red lines). The diagonal shows the density estimate for each variable.

## Discussion

The study of diversity on a regional scale (gamma diversity) shows the ability of environmental DNA to detect many taxa also identified by the traditional kicknet method. This result is in line with previous studies which reported several EPT taxa detected by both methods (Mächler et al. 2019, Seymour et al. 2021). However, a significant number of taxa known to be present in these rivers (according to the kicknet sampling) could not be detected by either the mlCOIintF/HCO2198 or fwhF2/EPTDr2n primers. In total, 23 EPT species were detected by kicknet and were not detected by either primer set. The non-congruence between kicknet and the eDNA methods is even more pronounced when results are assessed at local scale (alpha diversity). This result is not surprising, as pooling species information from multiple locations together across a region is likely to increase the set of species detected by both methods. It has been, however, a common practice in metabarcoding studies to perform comparisons at regional level (i.e. gamma diversity), which probably contributed to a misleading idea that eDNA and traditional methods are generally congruent. A recent meta-analysis showed, on the contrary, the low congruence between species list generated by DNA metabarcoding and traditional methods for macroinvertebrates (Keck et al. 2021). Thus, while numbers of diversity reported may be similar, the identity of taxa found by each method can substantially differ.

Overall, we observed a low correlation between the diversity measures estimated by the triad of different tested methods (kicknet, eDNA and model predictions). The highest correlation was found between eDNA (fwhF2/EPTDr2n primers) and the predictive model. This relationship might be to some degree driven by the fact that both methods reflect diversity at catchment scale as eDNA integrates to some point EPT diversity at the catchment level (Deiner et al. 2016) and the model estimates EPT diversity from multiple variables, catchment-wise (Kaelin and Altermatt 2016). The low correlations observed between the diversity measures estimated by the different methods can largely be explained by the methodological biases discussed above. It should also be noted that the locations studied have been sampled across a relatively limited gradient in river size (all were small to mid sized rivers), all between 370 to 912 m a.s.l. Therefore, the expected variation in the number of EPT species is limited and this reduces our ability to detect statistical relationships between the different methods. However, the variability in land-use in the catchments was relatively pronounced, such that arable land ranged between 0.1 and 81%, urban areas between 5 and 21%, and grassland between 4 and 54%. The main goal of our study, namely to use independent model predictions from a species distribution model (Kaelin & Altermatt 2016) to evaluate the accuracy of kicknet vs. eDNA approaches through a third, independent approach was only partially successful: indeed, the triad of approaches gave a triad of partially congruent and partially complementary results. The low congruence between the species detected by eDNA and kicknet can be explained by the numerous biases that can influence species detection probabilities at every step of data collection. For eDNA this can be caused by the complex dynamics of DNA in the environment (release rate by the organisms, degradation and dilution), manipulation of the DNA in the lab (conservation, extraction, PCR-amplification, sequencing), and the bioinformatics processing (Deiner et al. 2017). For the traditional methods, possible biases may concern sampling representativity (Larras and Usseglio-Polatera 2020) and taxonomic identification, including both errors and lack of precision (Stribling et al. 2008). However, the respective role of these factors remains difficult to disentangle and to estimate.

One of the reasons often cited to explain the non-detection of taxa by DNA methods is the incompleteness of reference databases (Weigand et al. 2019). This argument, although difficult to evaluate, is perfectly valid in studies dealing with the diversity of large or poorly known taxonomic groups (Lindeque et al. 2013). In the present study, this hypothesis can be excluded as all species detected by kicknet (except one) are present in the reference database used. However, this does not guarantee that the amplified regions can resolve all species detected by kicknet, nor that the intra-specific diversity of these species is fully represented in our reference database.

It should be noted that the choice of the primers and the barcode region to be amplified seems to play a significant role here. Overall, we found that fwhF2/EPTDr2n primers detected more EPT taxa than the mlCOIintF/HCO2198 primers. It appears that the taxa detected by the mlCOIintF/HCO2198 primers are in majority nested within the pool of taxa detected by the fwhF2/EPTDr2n primers, which is not surprising given that they are both amplifying a region of the same marker (COI). Hence our results confirm that for a group of organisms like the EPT, primer performance changes the detection rate on the exact same extracted eDNA sample. The fwhF2/EPTDr2n primers do have a higher target to non-target ratio for EPT compared to mlCOIintF/HCO2198 primers (but see Leese et al. 2021 for results and discussion for all benthic macroinvertebrates).

The fact that the more specific primers outperformed the less specific ones raises another important question: how many EPT species could not be correctly detected by the fwhF2/EPTDr2n primers because of their lack of specificity? It should be remembered that these primers, although more specific than the mlCOIintF/HCO2198 primers, cover a paraphyletic and very large group of organisms (basically, all insects, of which EPT make only a small percentage). Therefore, gains in the number of species detected by eDNA could be expected by using markers and primers specific to these three polyphyletic groups.

The large number of taxa detected only by the kicknet method should not mask the existence of several taxa that were detected only by their DNA. This result highlights the fact that DNA can provide real added value to traditional sampling techniques (Sweeney et al. 2011). The presence of these taxa can be explained on the one hand by the integrative aspect of environmental DNA, which reflects diversity on a larger scale via transport of DNA from upstream to downstream of the watershed (Deiner & Altermatt, 2014), and on the other hand by the capacity of DNA to identify species that are sometimes difficult to collect or identify using morphological criteria (Haase et al. 2006, Stribling et al. 2008).

In conclusion, our results suggest that the three approaches investigated here can give very different results about the species richness and the species composition of EPT communities. These differences are due to the respective biases of each method, but also to the different scales that they integrate. Kicknet sampling is carried out at one point and captures the organisms physically present at that location. In contrast, models typically provide estimates of macroinvertebrate diversity on a regular grid or at catchment level (Ferrier and Guisan 2006). Finally, environmental DNA is sampled at one point but has the characteristic of being transported from upstream to downstream, thus integrating diversity at the catchment scale (Deiner & Altermatt, 2014; Deiner et al. 2016). Therefore, although a certain degree of congruence is expected between the estimates produced by these methods, their different nature (observation vs. modelling) and the scales they incorporate can produce variable results, as shown here. Importantly, new frameworks integrating hydrological transport dynamics of eDNA allow to derive higher resolution diversity predictions and may act as a bridge between these methods (Carraro et al. 2020), yet have hitherto only been applied to catchments/scales larger than studied here. More efforts are needed to understand the reason why we observe such differences and additional work is needed to improve compatibility and comparability between them. However the achievable congruence between these approaches is currently limited as each comes with its own specificities, strengths and weaknesses. On the one hand, kicknet sampling and morphological identification and modeling are not likely to see major advancements that would change the outcome of our analysis. Whereas on the other hand, analysis of eDNA for macroinvertebrates still suffers from major drawbacks due to their paraphyletic origin and difficulty to exclude non-target groups during genetic analysis. Thus eDNA metabarcoding has the greatest potential for advancement through further method development and research. Here we showed that simply by changing primer sequences we could already improve correlation with the model. Regardless, until this challenge is solved, the three methods provide different perspectives on biological diversity and should be used together to provide complementary information to make informed decisions related to biodiversity management and conservation.

## Acknowledgements

We thank Marta Reyes for help during field work, Francis J. Burdon and Rik Eggen for comments on the project and coordination, and Pascal Stucki for sampling and identification of the EPT taxa. This work has been funded by the Swiss Federal Office for the Environment (BAFU/FOEN) and is part of the Eawag Ecoimpact initiative. Further funding is from The Swiss National Science Foundation (grant nr. <u>31003A_173074</u>), the University of Zurich Research Priority Programme in Global Change and Biodiversity (URPP GCB) to FA and the European Research Council (ERC) under the European Union’s Horizon 2020 research and innovation programme (Grant agreement No. 852621) to KD.

## Notes

### Competing Interest Statement

The authors have declared no competing interest.

