## Supplementary Information for "A triad of kicknet sampling, eDNA metabarcoding, and predictive modeling to assess aquatic macroinvertebrate biodiversity"

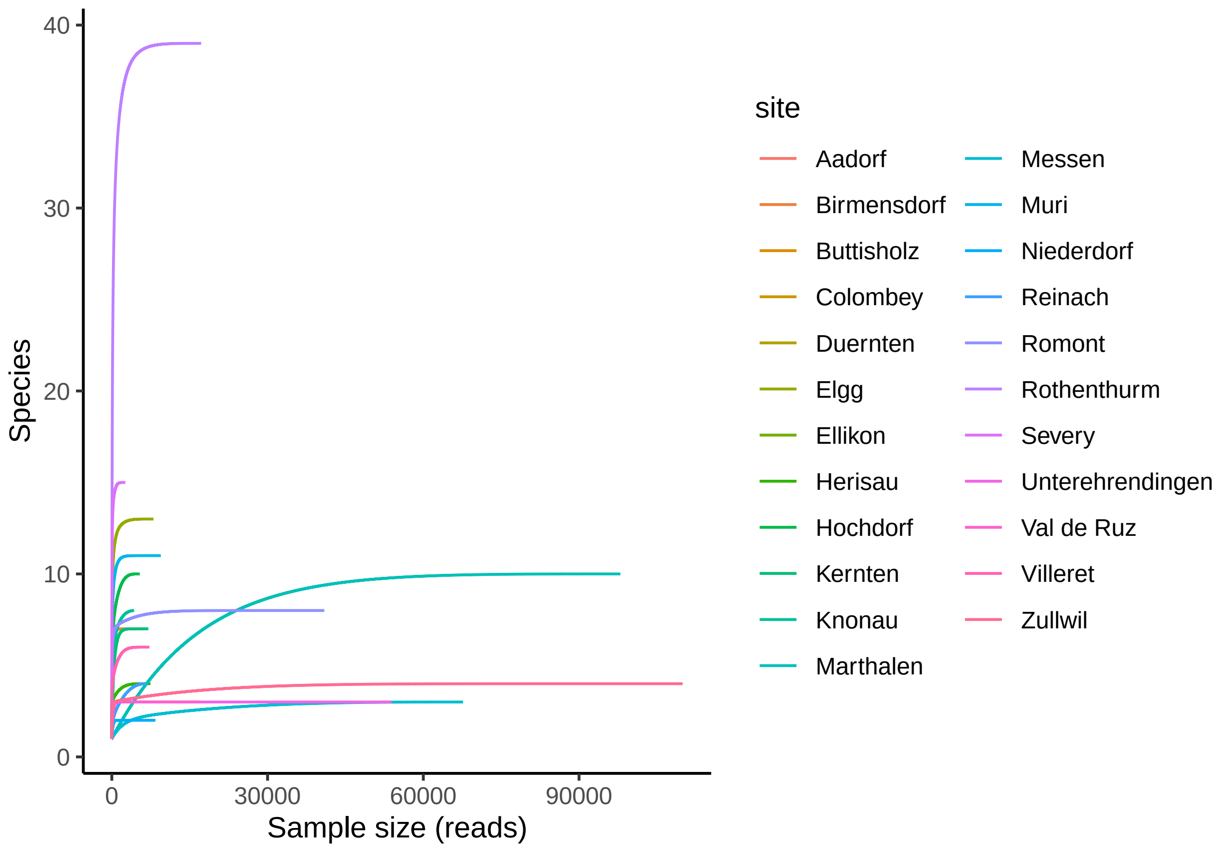

**Supplementary Figure 1.** Species accumulation curves of the EPT taxa detected using the mlCOI-intF/HCO2198 primers. Locations where EPT taxa were not detected are not shown.

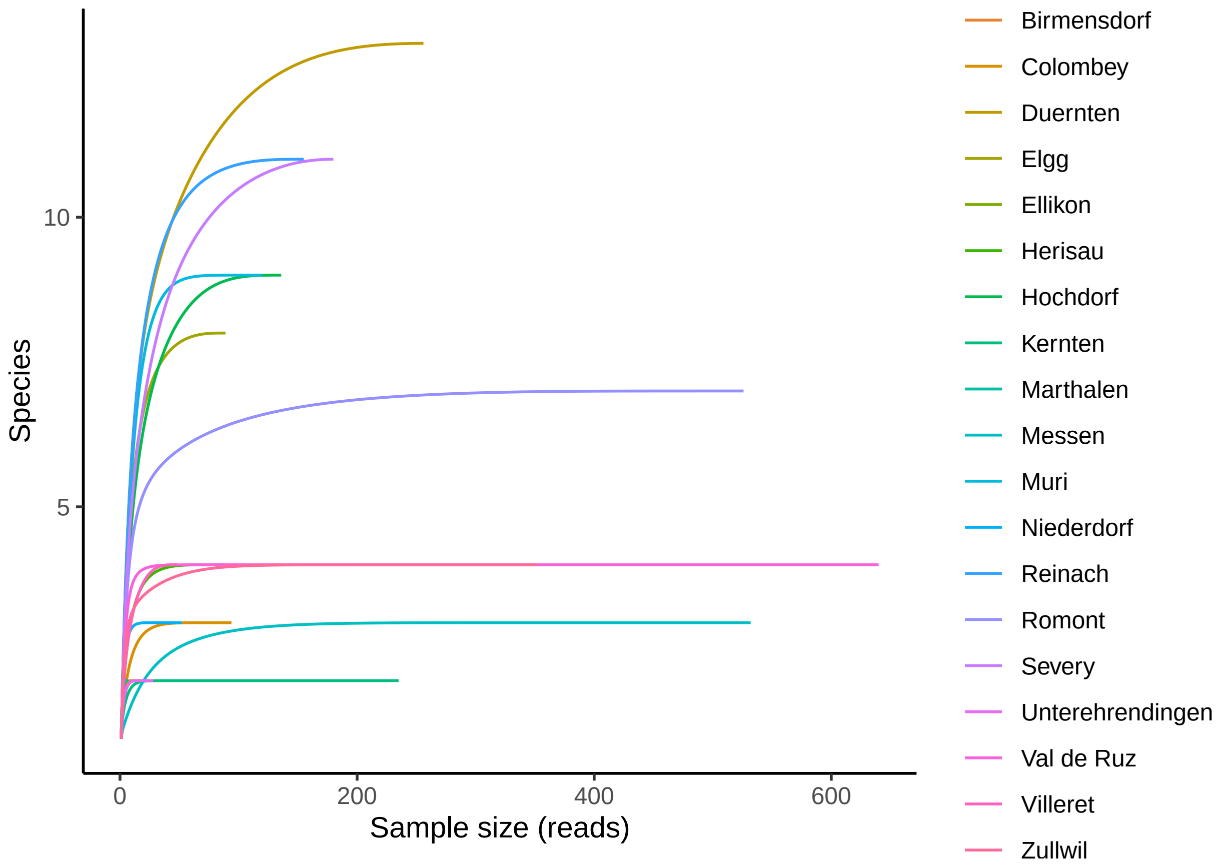

**Supplementary Figure 2.** Species accumulation curves of the EPT taxa detected using the fwhF2/EPTDr2n primers. Locations where EPT taxa were not detected are not shown.

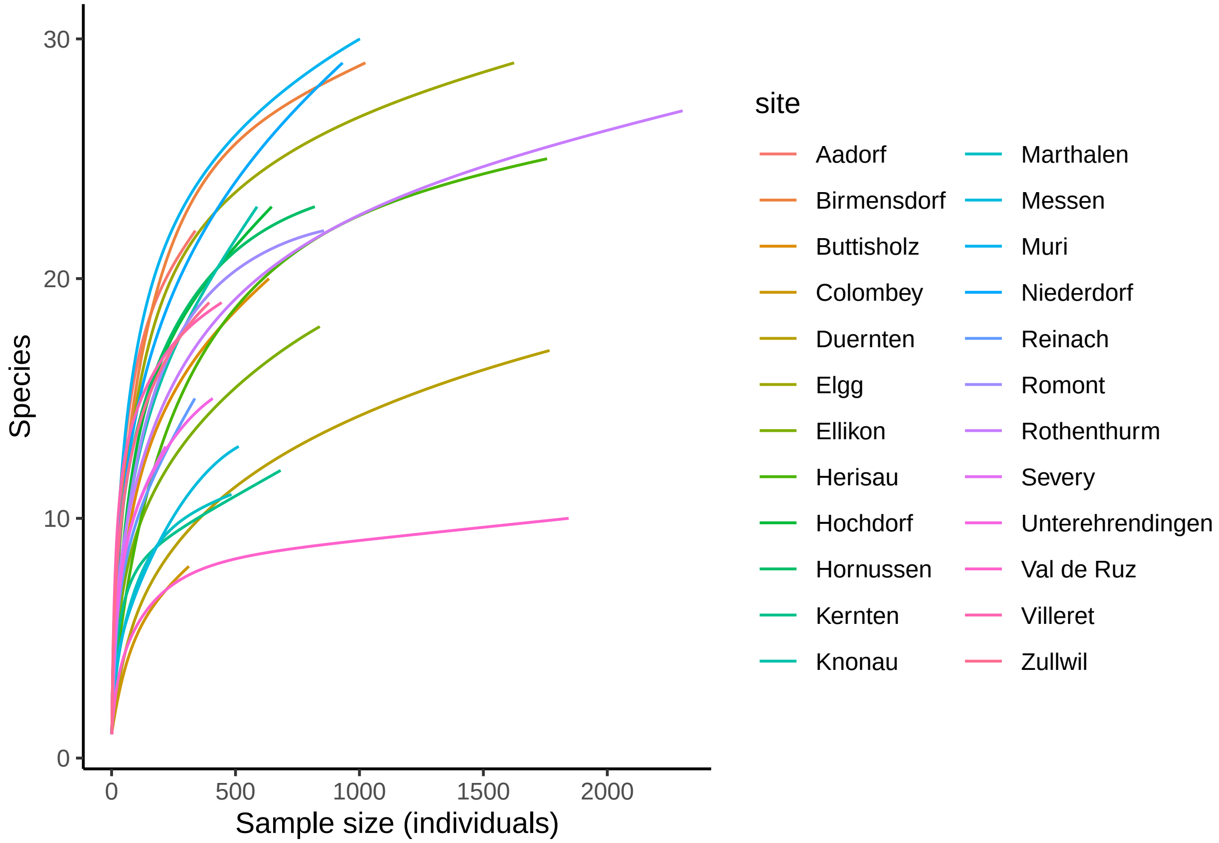

**Supplementary Figure 3.** Species accumulation curves of the EPT taxa detected using the kicknet method.

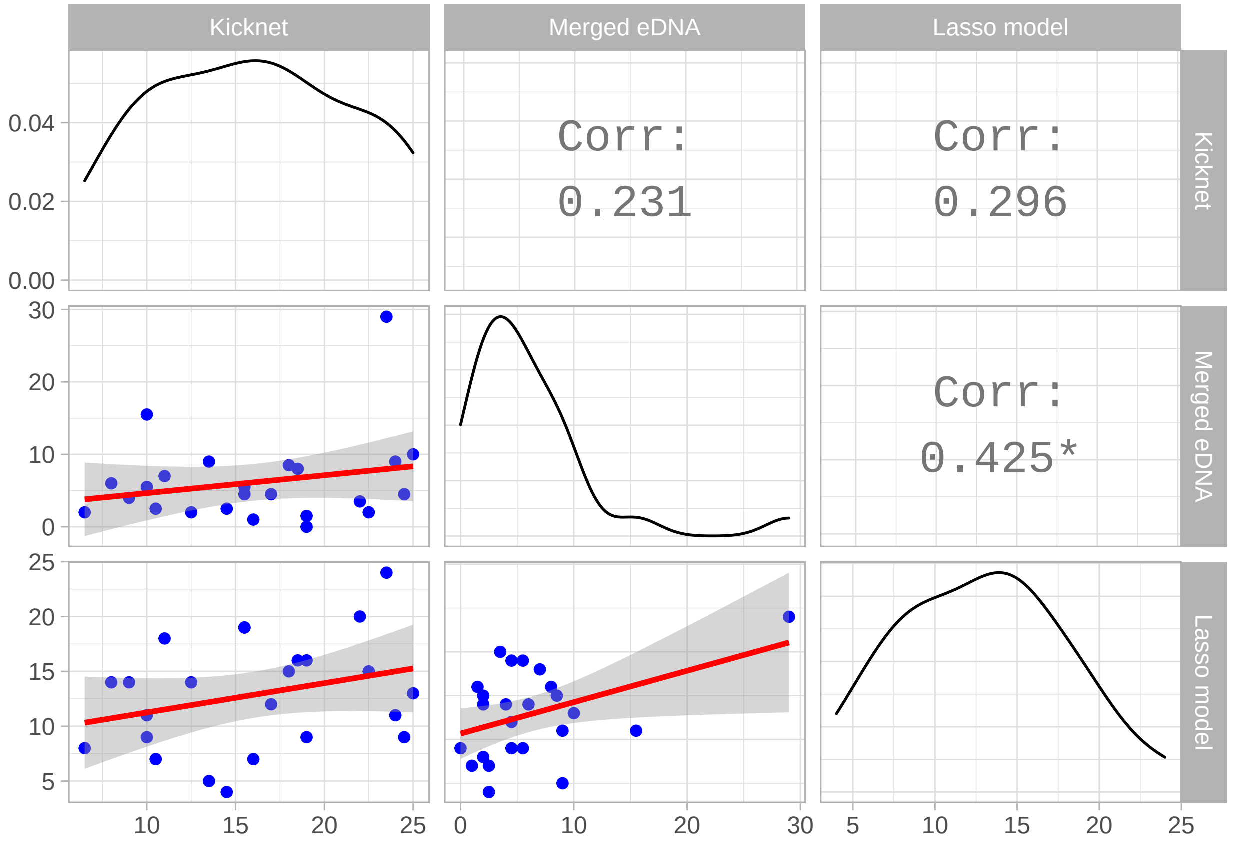

**Supplementary Figure 4.** Supplementary Figure 4. Relationships between the EPT richness estimates provided by the Kicknet, eDNA (mlCOIintF/HCO2198 and fwhF2/EPTDr2n primers merged), and the predictive model. The upper triangle provides the correlation values between each method (star indicates p-value $< 0.05$). Lower triangle shows the scatterplots with linear regressions (red lines). The diagonal shows the density estimate for each variable.
